# Harnessing the immune system to treat bone loss: The immunomodulatory and osteoprotective effects of the microalga *Skeletonema costatum*

**DOI:** 10.1101/2025.02.14.638280

**Authors:** Alessio Carletti, Katia Pes, Marco Tarasco, Joana T. Rosa, Sunil Poudel, Hugo Pereira, Bruno Louro, M. Leonor Cancela, Vincent Laizé, Paulo J. Gavaia

**Affiliations:** Centre for Marine Sciences, University of Algarve, Faro Portugal; Faculty of Medicine and Biomedical Sciences, University of Algarve, Faro, Portugal; S2AQUA - Collaborative Laboratory, Association for a Sustainable and Smart Aquaculture, Olhão, Portugal; GreenCoLab – Associação Oceano Verde, University of Algarve, Faro, Portugal; Algarve Biomedical Center, University of Algarve, Faro, Portugal

**Author notes:** Max Plank Institute for Hearth and Lung Research – Department of Developmental Genetics, Bad Nauheim, Germany (Alessio Carletti, Marco Tarasco). Center for Transplantation Sciences, Massachusetts General Hospital, Harvard Medical School, Boston (Sunil Poudel). Excellence Cluster Cardio-Pulmonary Institute CPI, Justus-Liebig University of Giessen, Giessen, Germany. Correspondence: Paulo J. Gavaia (.).

**Keywords:** Osteoimmunology, Inflammation, T-cell activation, Antigen presentation, Microalgae, Immunoporosis

## Abstract

The emerging field of osteoimmunology provides compelling evidence for the pivotal role of the immune system in the development of bone erosive pathologies such as osteoporosis. However, no immunomodulatory drug has yet been integrated into the therapeutic management of bone loss. Recently, driven by the demand for next-generation treatments for these conditions, natural compounds are gaining renewed attention as promising candidates for drug discovery.

In this study, we explored the anti-osteoclastogenic effects of an emerging extract from the marine microalga *Skeletonema costatum*. Using a zebrafish model of bone regeneration, we demonstrated the extract’s ability to inhibit the recruitment of osteoclast progenitors and block their differentiation into mature osteoclasts *in vivo*. Bulk RNA sequencing of early-stage fin blastemas revealed the downregulation of genes involved in inflammation, T-cell activation, and antigen presentation, suggesting that the extract exerts its effects primarily through immunomodulatory mechanisms. To further assess its therapeutic potential, we tested the extract in a medaka model of RANKL-induced osteoporosis and on a murine macrophage cell line. The extract effectively prevented bone loss in fish and inhibited osteoclastic differentiation in murine macrophages *in vitro*.

Collectively, our findings provide mechanistic insights into a novel, therapeutically relevant natural extract, offering proof of concept for its osteoprotective potential through immune system modulation.

**Significance:** Recent findings in the field of osteoimmunology reveal the potential of targeting immune cells to regulate bone homeostasis. However, this approach has yet to be applied to therapies for bone erosive conditions. This study explores the potential of an immunomodulatory strategy using an emerging natural extract, which prevent osteoclast differentiation by modulating inflammation, T-cell activation, and macrophage fate determination in zebrafish and medaka models of bone regeneration and osteoporosis. The extract also inhibits osteoclastic differentiation in a murine macrophage line, suggesting its translatability to mammalian systems. By focusing on immune pathways, this research provides a proof of concept for developing immunomodulatory treatments for osteoporosis and similar conditions, addressing a critical need in bone health management.

## Introduction

Bone erosive pathologies such as osteoporosis are an urgent medical challenge that require immediate attention^1,2^. They are metabolic bone disorders linked to the disruption of the equilibrium between bone formation and bone resorption, leading to mineral loss and bone structural deterioration^3,4^. Therapeutically, these diseases are managed with a relatively small repertoire of pharmaceuticals, which typically, directly target bone-forming cells (osteoblasts), aiming at increasing the endogenous osteogenesis, or bone-resorbing cells (osteoclasts), by inhibiting their differentiation, action, or life span.

Examples of osteoanabolic drugs are parathyroid hormone analogues (e.g. teriparatide and abaloparatide)^5–7^, while anti-resorptive agents include bisphosphonates and RANKL-monoclonal antibodies (e.g. Denosumab)^8–10^. Recently, dual action drugs like the anti-sclerostin monoclonal antibody Romosozumab have proven to be a promising option for the treatment of bone erosive disorders^11^. These molecules have shown good efficacy in the short-term, and currently represent the spearhead of therapy for various bone erosive disorders, but are not exempt from limitations. All options tend to lose efficacy over time^12^, and while antiresorptive drugs are associated with rare yet serious complications, such as osteonecrosis of the jaw^13,14^, bone anabolic drugs have been associated with the occurrence of osteosarcoma^5–7^.

The limited provision of therapeutics to treat bone loss has generated a strong demand for the development of new approaches. This need is driving deeper investigations into the mechanisms behind bone erosive disorders and encouraging the pharmaceutical exploration of novel osteoactive molecules that offer long-lasting effects without side effects.

In this regard, the key role of the immune system in regulating bone metabolism is supported by a large body of research, and osteoimmunology has recently gained in popularity as a new research field^15^. Immune cells exert a tightly regulated control over bone homeostasis^16,17^.

Among these, macrophages share a common monocytic lineage with osteoclasts, which sparked the definition of osteoclast as bone macrophages^18^, and represent a major source of osteoclasts in vertebrates including fish, mice and humans^19,20^. Together with macrophages, T-cells are the main cellular regulators of inflammation, a potent inducer of bone resorption. Inflammation is required for the receptor activator of the nuclear factor kappa-Β ligand (RANKL) signaling pathway, the central molecular driver of osteoclast differentiation^21–25^. Chronic inflammation is widely recognized as a key root cause of bone loss in various erosive pathologies^26^. For example, postmenopausal osteoporosis is increasingly acknowledged as an inflammatory disease, and was (re)named immunoporosis^27–30^, where activated T-cells establish a chronically increased bone resorption by stimulating osteoclast differentiation through RANKL-dependent and independent pathways^31,32^. The leading role of the immune system in the pathophysiology of osteoporosis and other bone disorders raises the question of whether immunomodulatory therapies could effectively prevent bone loss in patients suffering from such disorders. Surprisingly, little investigation has been conducted in this regard so far on the potential of targeting immune cells to control bone loss.

The search for novel therapeutic approaches has recently refreshed the role of natural compounds, with marine-derived molecules becoming promising assets in drug discovery^33,34^.

This is particularly true for metabolic bone disorders, as an increasing number of studies have identified molecules and extracts obtained from marine organisms with immunomodulatory and osteoprotective bioactivities^35^. Among those, microalgae are attracting a significant interest, due to a combination of their capacity to synthesize bioactive compounds, technologically advanced cultivation technologies^36–38^, and their amenability to genetic engineering^39,40^.

Here, we have explored, as a proof of concept for the potential of natural immunomodulatory molecules for the treatment of bone loss, the bioactivity of the microalga *Skeletonema costatum*, a marine species commonly cultivated in Europe and approved for human consumption^41^. The ethanolic fraction from this microalga have recently shown a potent anti-inflammatory activity *in vitro*^42,43^. Here we have characterized the anti-osteoclastogenic activity of the *S. costatum* ethanolic extract by taking advantage of a zebrafish model of bone regeneration^44^, and investigated the underlying molecular mechanisms by tissue-specific transcriptomics. Subsequently, we have evaluated the anti-osteoporotic potential of the algae in a medaka genetic model of osteoporosis^45^, and validated its capacity to prevent osteoclastic differentiation using a murine macrophage cell line.

## Results

### Skeletonema suppresses osteoclast recruitment during caudal fin regeneration

The impact of the ethanolic fraction of *Skeletonema costatum* (SKLT, hereafter) on bone regeneration was first assessed using the zebrafish caudal fin assay. The system was chosen as it provides a relatively simple and standardized method to investigate the concerted interaction of different cellular types in the *de novo* formation of bone structures, and is particularly suitable for in vivo cell-tracking^44,46^. Following caudal fin amputation, adult zebrafish were exposed to SKLT via immersion in water supplemented with the extract. Regenerative and mineralogenic performances and morphogenic patterning of regenerating bony rays were determined at a single time-point, 5 days post-amputation (dpa, **Figure 1A**, **B**). Regenerated area in SKLT-treated fish showed a tendency to a reduction although it was not significant (**Figure 1C**). However, SKLT-exposed animals showed a clear increase in the mineralized area (**Figure 1D**). SKLT also affected the patterning of the bony rays. While ray width was not affected (**Figure 1E**), ray bifurcation suffered a distalization along the proximal-distal axis in SKLT-treated fish (**Figure 1F**). These observations suggest that SKLT exposure affected the pattern of mineral deposition, indicating a potential switch in the equilibrium between bone formation and resorption^47^.

**Figure 1.**
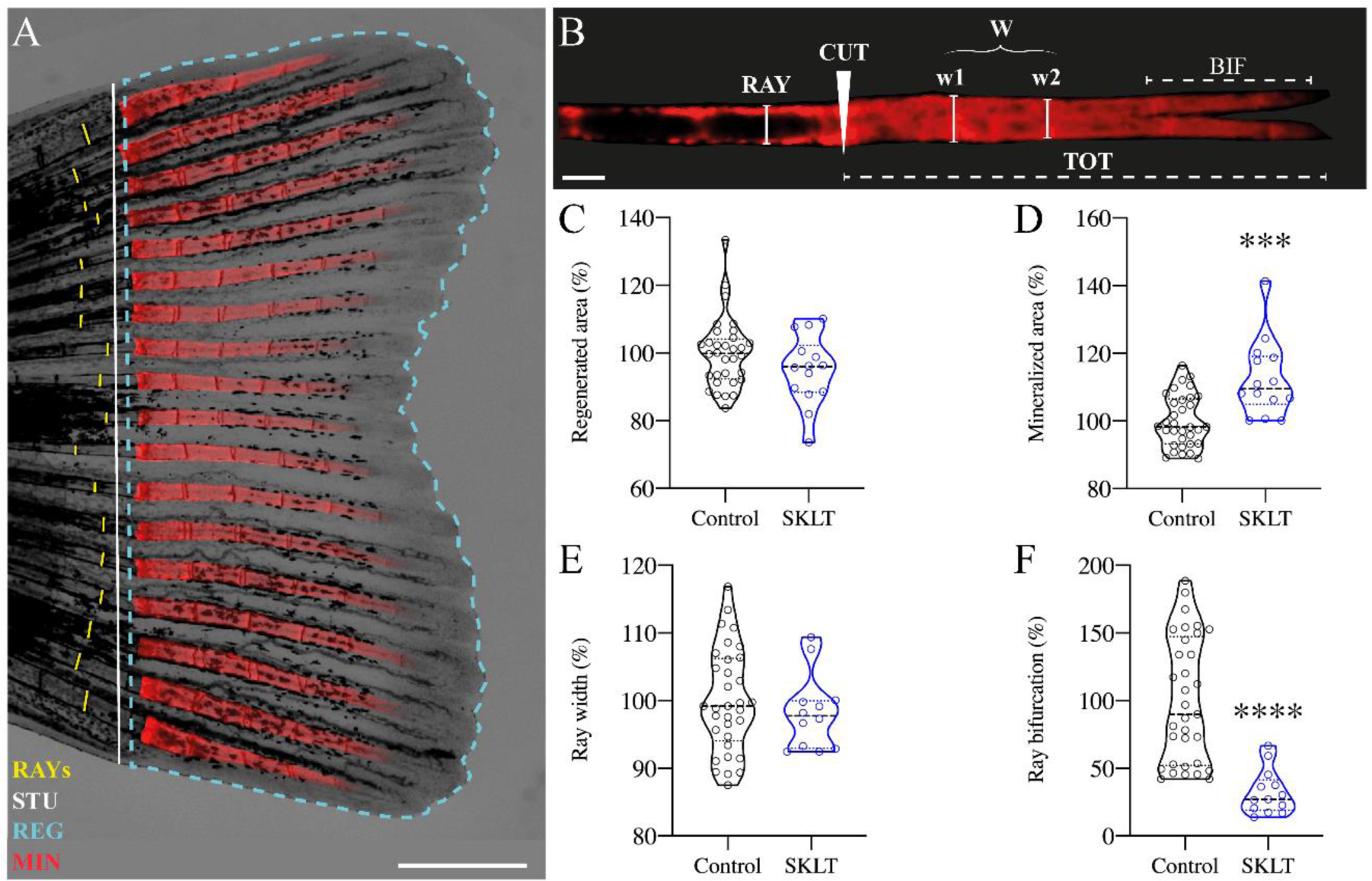
*Skeletonema costatum* extract shifts the mineral equilibrium of regenerating rays towards pro-mineralogenic outcomes. Caudal fin regeneration and mineralization were assessed in young adults exposed to the ethanolic extract of *S. costatum* (SKLT) or 0.1% ethanol (Control). (**A**) Representative image of a caudal fin illustrating the morphometric measurements used to assess regenerative and mineralogenic effects of SKLT, i.e. ray width (RAY), stump length (STU), regenerated area (REG) and mineralized area (MIN). (**B**) Representative image of an alizarin red S (AR-S) stained fin ray illustrating the morphometric measurements used to assess the patterning effect of SKLT, i.e. amputation plan (CUT), ray width before the amputation (RAY), average ray width calculated as the average width of the first two segments after the amputation plan (W), length of bifurcation (BIF) and total length of regenerated ray (TOT). (**C-F**) Effect of SKLT on the regenerated area (**C**), mineralized area (**D**), average ray width (**E**) and ray bifurcation (**F**). Normality was tested through a Anderson-Darling test (*p*<0.05). Statistical differences were tested through an unpaired *t* test (*p*<0.05) or a non-parametric Mann-Whitney test (*p*<0.05) whenever the data distribution resulted non-normal. Asterisks indicate values statistically different with *p*<0.0005 (***) and *p*<0.0001 (****). Scale bars are 1 mm in **A** and 250 µm in **B**.

We questioned whether there was involvement of an anti-resorptive mechanism underlying this effect, and decided to monitor the dynamics of osteoclast precursors, identified as rounded cathepsin k (*ctsk*)-expressing cells, in Tg(Ola.*ctsk*:FRT-DsRed-FRT-Cre,*myl7*:EGFP)^mh201^ transgenic zebrafish^48^, hereafter referred to as Tg(*ctsk*:DsRed), over 10 days of continuous exposure following caudal fin amputation (**Figure 2A**). In vehicle-treated fish (CTRL), the population of *ctsk^+^* cells peaked between 24 and 48 hours post-amputation (hpa) (**Figure 2B**, **D**), then their density in the regenerated area declined over time. In contrast, significantly fewer *ctsk^+^* cells were recruited to the blastema in SKLT-treated fish at 24 and 48 hpa (**Figure 2C**, **D**). Following a peak at 72 hpa, the number of *ctsk^+^* cells was similar in SKLT-treated and control fish. Elongated tubular *ctsk*-expressing osteoclasts resembling osteolytic tubules (OLTs) recently described^49^, were observed beside the regenerated rays as early as 96 hpa in control fish and 120 hpa in SKLT-treated fish (**Supplementary Figure S1**). The reduction in the number of *ctsk^+^* cells was associated with a decrease in resorptive activity demonstrated by the almost complete suppression of TRAP staining at 24 hpa, but not at 240 hpa (**Figure 3A**, **B**). This suggests that SKLT may have the capacity to inhibit the recruitment of osteoclast precursors in the early stages of fin regeneration, although not sufficiently to fully suppress their differentiation into functional osteoclasts later on. Interestingly, in the second experiment (10 days of exposure), SKLT-treated fish showed a significant reduction of the regenerated area starting from 72 hpa until 240 hpa (**Figure 4**), suggesting that the extract might affect molecular pathways involved in the expansion of the regenerating blastema.

**Figure 2.**
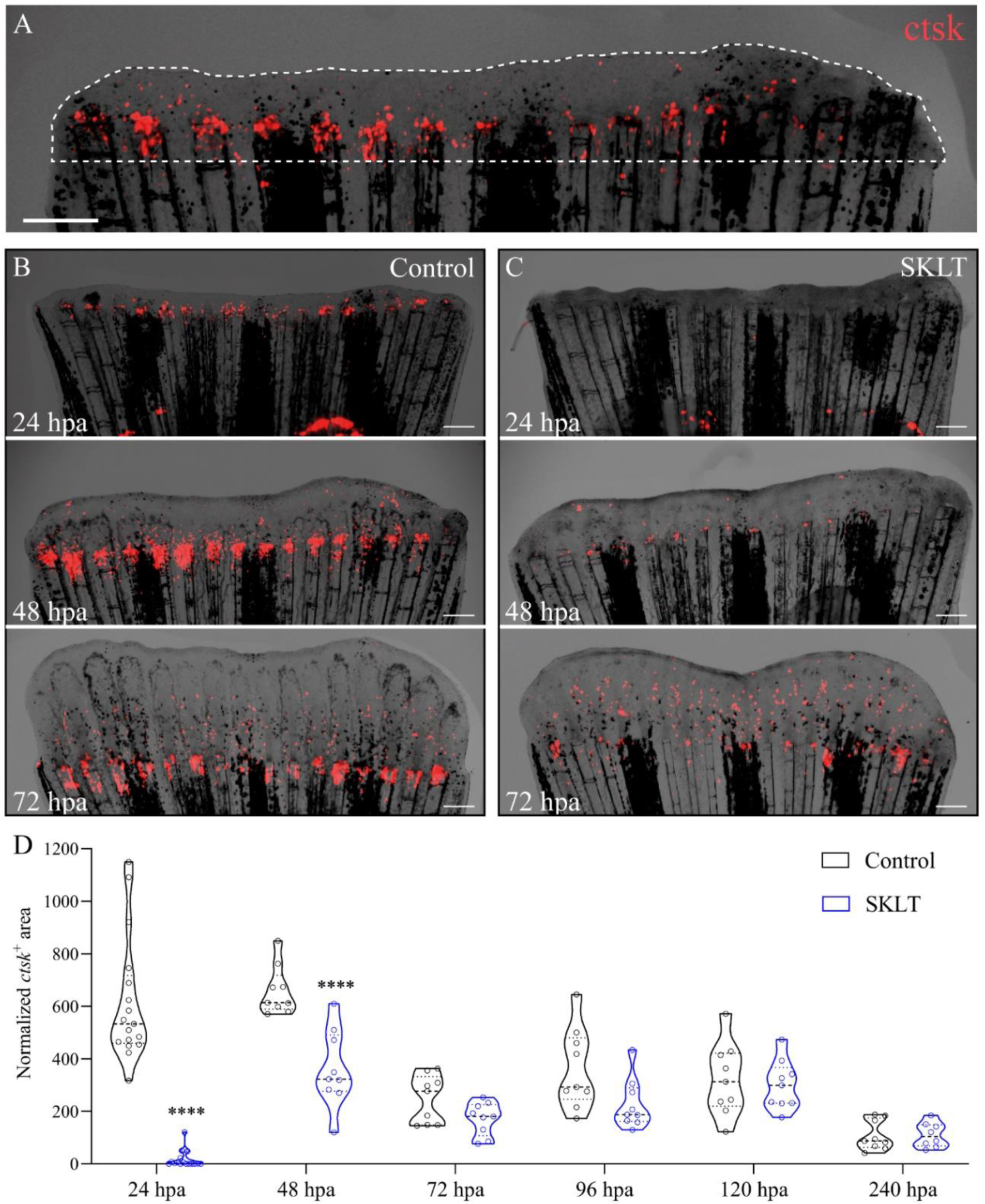
SKLT inhibits the recruitment of osteoclast precursors to the caudal fin blastema. Osteoclastic recruitment was determined from DsRed fluorescence signals observed in the regenerating caudal fin of transgenic fish Tg(*ctsk*:DsRed) (**A**) Representative image illustrating the regenerated area (dotted line) and *ctsk^+^* area (ctsk, red signal). Time-course of *ctsk*^+^ area in regenerating fins of SKLT-treated (**B**) and control fish (**C**). (**D**) Quantification of *ctsk*^+^ area in regenerating fins of SKLT-treated and control fish from 24 to 240 hours post-amputation (hpa). At each timepoint, statistical differences were tested through and unpaired *t* test (*p*<0.05) or a non-parametric Mann-Whitney test (*p*<0.05) whenever the data distribution resulted non-normal. Asterisks indicate values statistically different with *p*<0.0001 (****). Scale bars are 500 µm.

**Figure 3.**
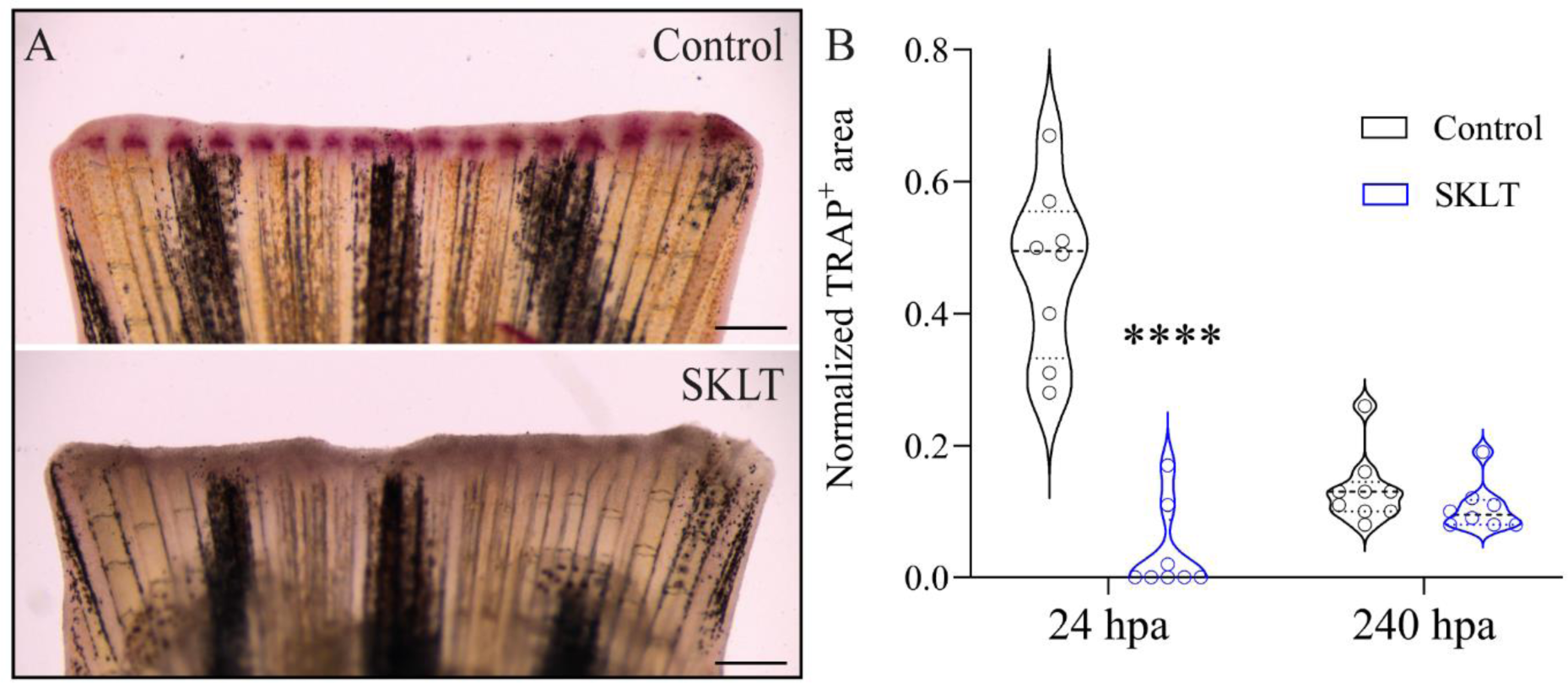
SKLT suppresses bone resorption activity at 24 hours post-amputation. Osteoclast activity was assessed through TRAP (tartrate-resistant acid phosphatase) staining in the regenerating caudal fin of adult zebrafish exposed to the ethanolic extract of *S. costatum* (SKLT) or 0.1% ethanol (Control) at 24 and 240 hours post-amputation (hpa). (**A**) Representative images showing TRAP staining at 24 hpa in control and SKLT-treated fish. **(B)** Quantification of TRAP^+^ area at early regeneration (24 hpa) and late regeneration (240 hpa) stages. Statistical differences were tested through an unpaired *t* test (*p*<0.05). Asterisks indicate values statistically different with *p*<0.0001 (****). Scale bars are 500 µm.

**Figure 4.**
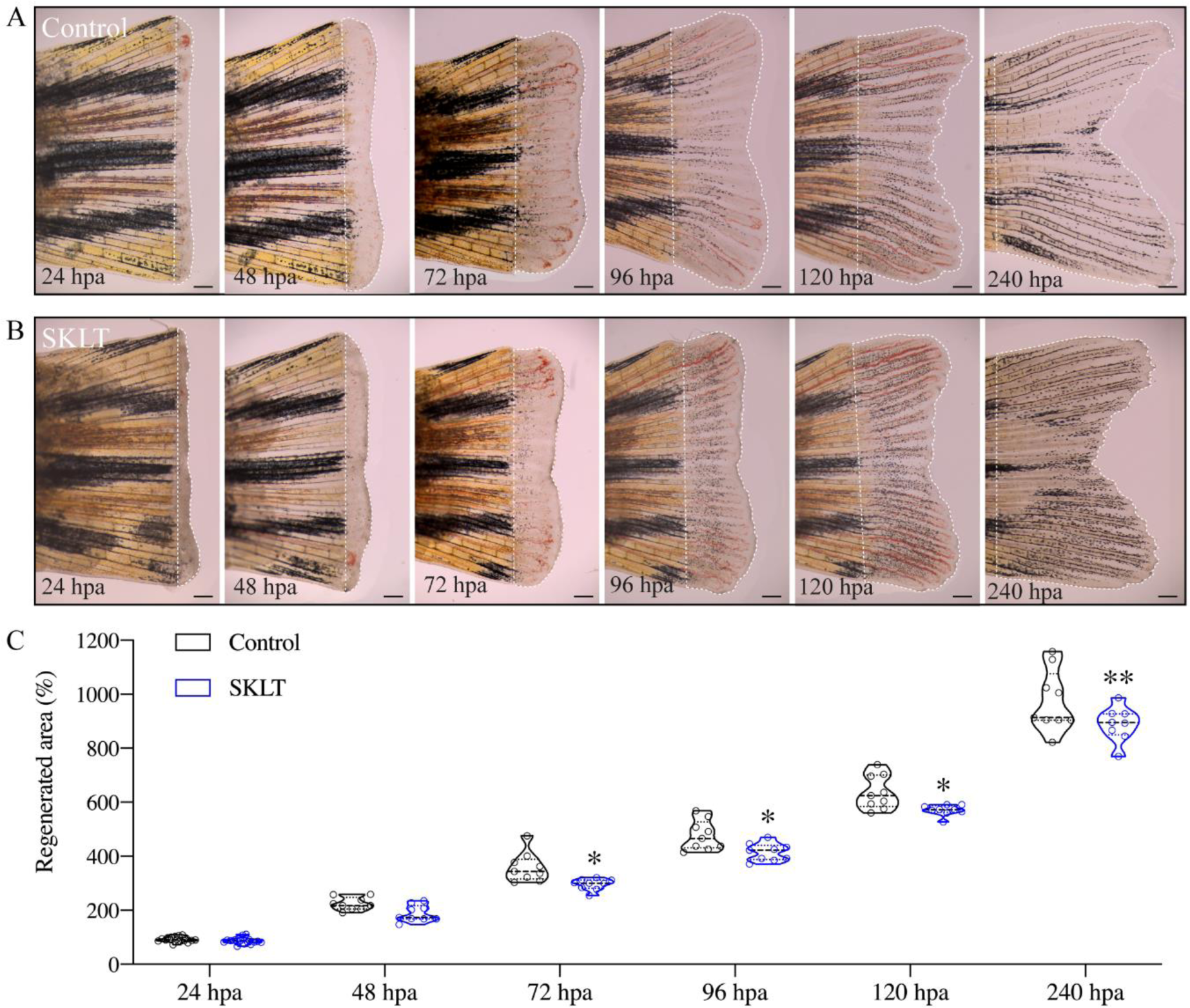
SKLT reduces the overall regenerative performance. Regeneration of the caudal fin was assessed in adult zebrafish exposed to the ethanolic extract of *S. costatum* (SKLT) or 0.1% ethanol (Control) from 24 to 240 hours post amputation (hpa). (**A-B**) Representative images depicting the time-course of caudal fin regeneration (dotted line indicates regenerate) in control (**A**) and SKLT-treated (**B**) fish. (**C**) Quantification of the regenerated area at relevant time points. Statistical differences were tested through an unpaired *t* test (*p*<0.05) or a non-parametric Mann-Whitney test (*p*<0.05) whenever the data distribution resulted non-normal. Asterisks indicate values statistically different with *p*<0.05 (*), *p*<0.01 (**). Scale bars are 500 µm. Picture illumination was artificially adjusted for better visualization.

### SKLT modulates immune processes and suppresses inflammation, T-cell activation, and antigen presentation in the regenerating fin blastema

Because SKLT was previously shown to possess anti-inflammatory activity *in vitro*^42,43^, and given the importance of inflammation in activating pro-regenerative molecular programs^50^, we hypothesized the existence of an immunomodulatory mechanism behind the effect observed in SKLT-treated fish. We investigated the molecular mechanisms underlying SKLT inhibition of pre-osteoclast recruitment through tissue-specific transcriptomics, by dissecting the early regenerated blastema at 24 hpa, corresponding to the time-point at which we observed the strongest suppressive effect over the recruitment of osteoclasts progenitors.

Bulk RNA-Seq data identified 720 differentially regulated genes (FDR < 0.01). Of those, 361 were upregulated, and 359 were downregulated (**Supplementary Figure S2**). Differentially expressed genes were assigned gene ontology using the online platform DAVID, and gene list analysis for Biological Processes (BP) evidenced 165 significantly enriched terms. BP terms were filtered for uniqueness and dispensability to compensate for redundancy, and resulting BPs were plotted with REVIGO (**Figure 5A**). Subsequently, we prepared gene expression data heat maps for the most relevant enriched biological processes.

**Figure 5.**
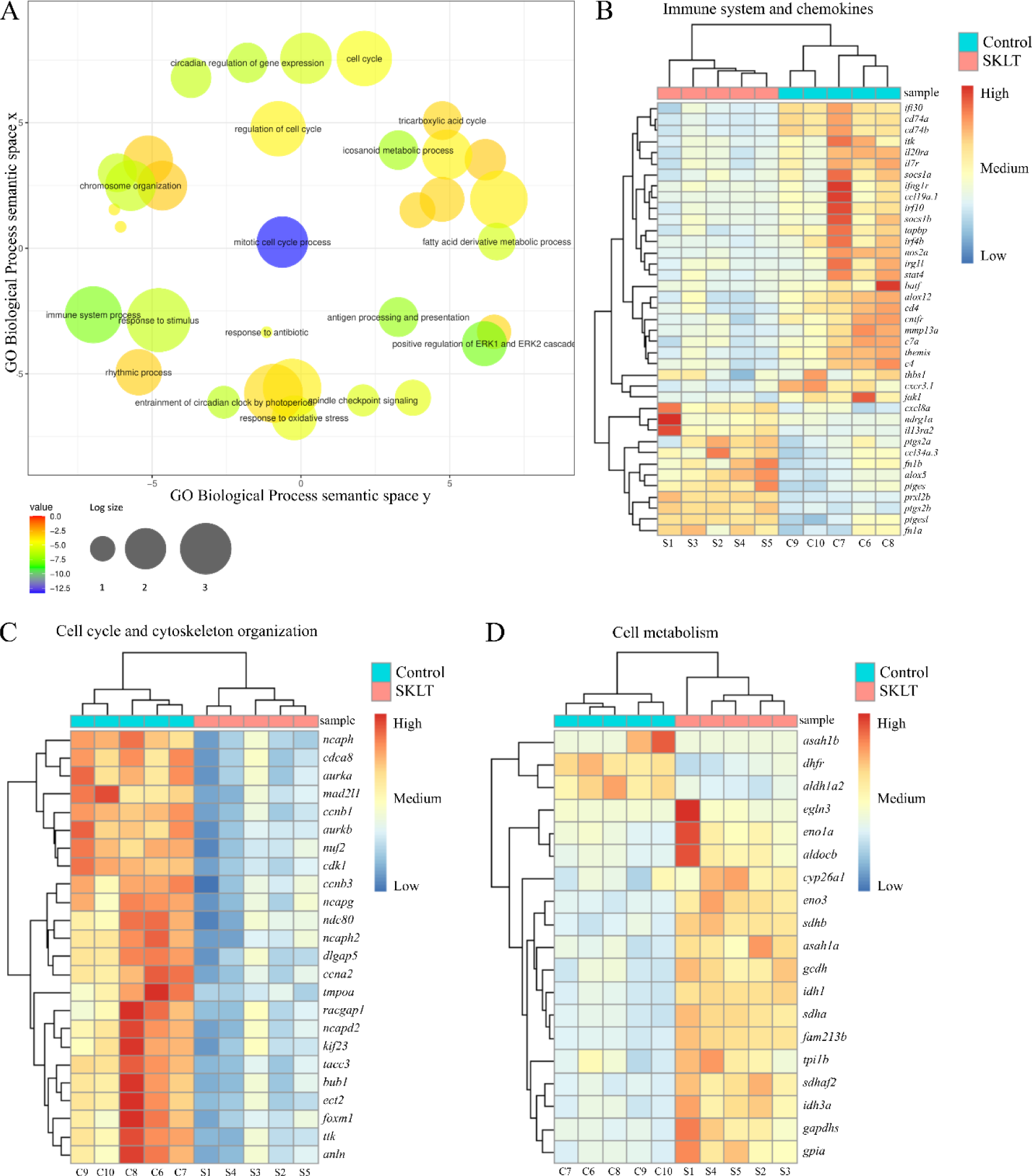
Analysis of the transcriptome of regenerating caudal fin blastemas reveals that SKLT modulates immune response, cell cycle and metabolism. RNA-Seq analysis identified differentially expressed genes (DEGs) and enriched gene ontology terms for caudal fin blastemas at 24 hours post-amputation (hpa) in control and SKLT-exposed fish. (**A**) REVIGO plot of biological processes filtered for redundancy and dispensability. Cluster analysis and heat map of differentially expressed genes associated with (**B**) Immune system (GO:0002376), Eicosanoid metabolism (GO:0006690), Inflammatory response (GO:0006954), T cell activation (GO:0042110), and Cytokine-mediated signaling pathway (GO:0019221). (**C**) cell cycle (GO:0022402) and cytoskeleton organization (GO:0007010); (**D**) glycolysis (GO:0006096), fatty acids metabolism (GO:0006099), tricarboxylic acid cycle (GO:0006631), and carboxylic acid metabolic process (GO:0019752).

Among the most regulated biological processes there were genes linked to adaptive and innate immune responses. In detail, genes involved in inflammation (*irg1l*, *nos2a*, *thbs1*, *il20ra*), the interferon system (*ifi30*, *ifng1r*, *irf4b*, *irf10*), and the complement system (*c4*, *c7a*), were downregulated in blastemas of SKLT versus control fish (**Figure 5B**). Interferon γ (IFN-γ), which is mainly produced by Th1 cells^51^, induces antigen-dependent T cell activation, and lead to the production of pro-osteoclastogenic inflammatory cytokines^51^. Interestingly, genes involved in T-cell activation and antigen presentation (*socs1a*, *itk*, *il7r*, *batf*, *cd4*, *cd74a*, *cd74b*, *jak1*)^31^, were also downregulated in SKLT-exposed fish (**Figure 5B**). Conversely, a second cluster of immunoregulatory genes was upregulated upon SKLT exposure, including fibrins (*fn1a*, *fn1b*) and interleukin 13 receptor alpha 2 (*il13ra2*). The expression of genes involved in paracrine communication through chemokines was regulated in a dualistic manner. While genes encoding for chemokines Ccl34 (*ccl34a.3*) and Cxcl8 (*cxcl8a*) were upregulated, the genes encoding for Ccl19 (*ccl19a.1*), and the receptor Cxcr3 (*cxcr3.1*) were downregulated. Overall, these data support the hypothesis that *S. costatum* induces a multifaceted immunomodulation on regenerating blastemas, which ultimately leads to reduced osteoclast formation.

### SKLT inhibits retinoic acid production and cell proliferation, and stimulate energy production in the regenerating fin blastema

Cluster analysis of genes involved in cell cycle control and cytoskeleton organization revealed that exposure to SKLT may suppress cell proliferation in 24 hpa blastemas (**Figure 5C**). Among downregulated genes, those involved in cell proliferation (*ttk*, *cdca8*, *dlgap5*, *tacc3*, *nuf2*, *ndc80*), mitosis (*bub1*, *ccnb1*, *ccnb3*, *ccna2*, *cdk1*, *tmpoa*, *mad2l1*, *foxm1*) and cytokinesis (*ect2*, *aurka*, *aurkb*, *kif23*, *racgap1*, *anln*) were the most impacted by SKLT. A deeper look at genes involved in energy metabolism (**Figure 5D**) revealed a significant up-regulation of genes involved in glycolysis (*eno3*, *eno1a*, *gapdhs*, *aldocb*, *tpi1b*, *gpia*) and fatty acids oxidation (*gcdh*, *asah1a*, *sdhb*, *sdhaf2*, *idh1*, *sdha*, *idh3a*) upon exposure to SKLT. Interestingly, *aldh1a2* (coding for Aldehyde dehydrogenase 1 family, member a2) was downregulated in SKLT-treated fish. Aldh1a2 is responsible for the production of retinoic acid (RA), and gene expression is highly upregulated at early stages of fin regeneration^52^. Because RA drives proliferation and growth of the blastema^52^, and exerts a dynamic control over osteoblast dedifferentiation, proliferation, and redifferentiation during fin regeneration^53^, a reduction of RA synthesis is coherent with the inhibition of genes involved in cell proliferation (**Figure 5C**) in early stage regenerating blastemas (24 hpa), and the decreased regenerative outgrowth observed at later stages in fish treated with SKLT further supports this model (**Figure 4**).

### SKLT protects from bone loss in a medaka model of osteoporosis

Following the results obtained in zebrafish, we questioned whether the immunomodulatory properties of SKLT could prevent bone loss in a medaka inducible osteoporosis model, Tg(*rankl*:HSE:CFP)^TG1135^, where bone loss is triggered by conditionally overexpressing RANKL^19^. Successful induction is tracked by monitoring the fluorescence of CFP (cyan fluorescent protein), whose co-expression with RANKL is driven by a bidirectional heat-shock promoter. A higher CFP signal is indicative of a higher RANKL expression thus of a higher resorptive activity and induction of bone loss. RANKL expression was estimated by measuring CFP fluorescence intensity in heat-shocked medaka larvae exposed for 6 days to SKLT or its vehicle (**Figure 6A**, **B**). Because frequency distribution of CPF fluorescence intensity was bimodal or right-skewed, we decided to cluster fish into 2 subgroups, based on their CFP fluorescence intensity (**Figure 6C**). We then determined the induction of bone loss by measuring the mineralized area of the vertebral column by alizarin-red staining. No differences were observed between control and SKLT in the groups showing low levels of CFP expression, which proxies a low RANKL expression (**Figure 6D**). On the contrary, in the group that included fish with high CFP/RANKL expression, the level of mineralization of the vertebral column was significantly increased upon exposure to SKLT (**Figure 6E**). Those fish also displayed a higher number of mineralized neural arches (**Figure 6F-H**). Such observation indicates that while no protective effect of SKLT over bone loss was present in fish suffering a mild RANKL induction, bone loss was significantly reduced upon exposure to SKLT in fish with the strongest RANKL induction, and therefore, fish were protected from developing an osteoporotic phenotype.

**Figure 6.**
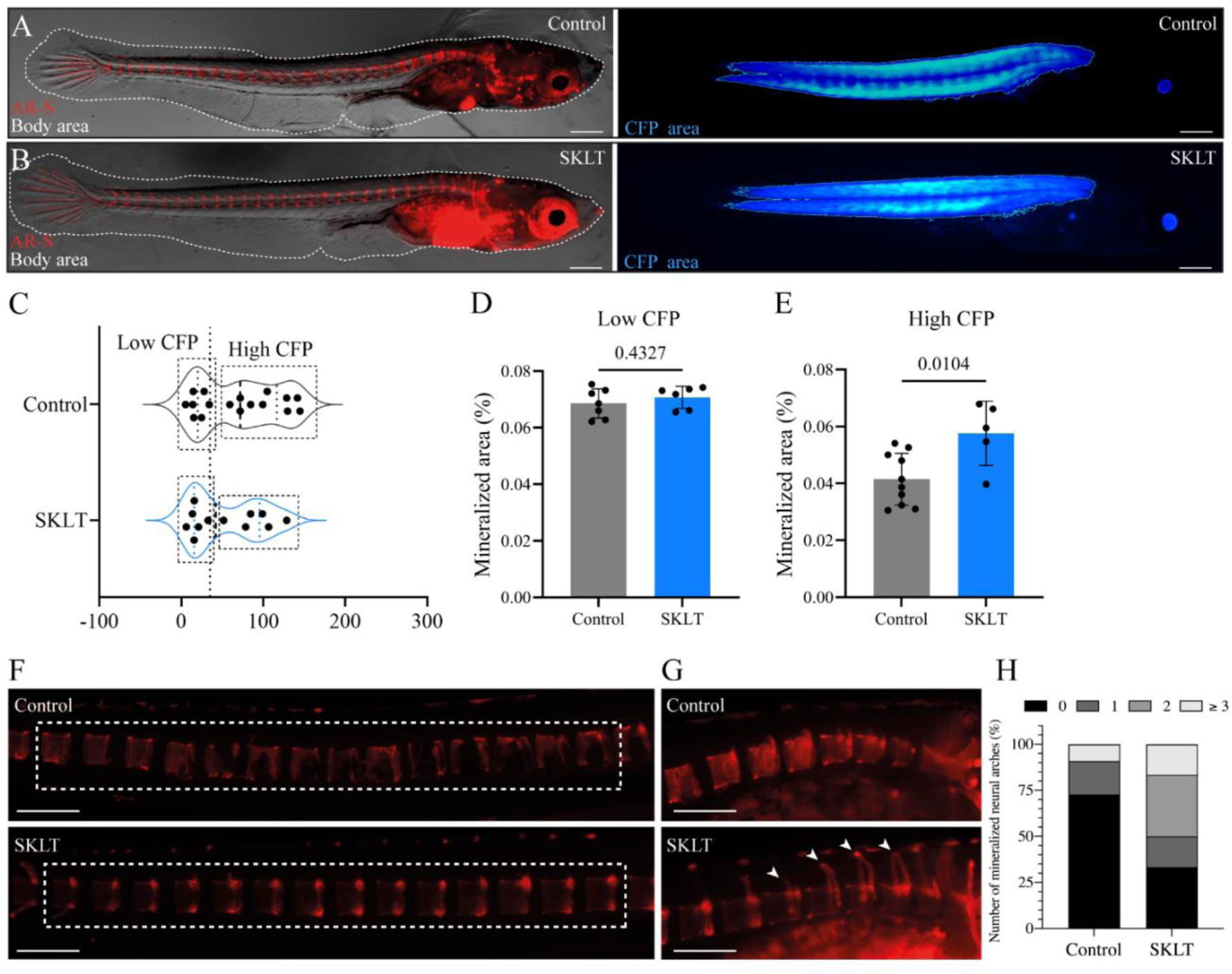
SKLT protects from bone loss in a medaka model of RANKL-induced osteoporosis. Mineralization of the vertebral column was assessed in adult medaka induced for osteoporosis and exposed to the ethanolic extract of *S. costatum* (SKLT) or 0.1% ethanol (Control). (**A-B**) Representative images illustrating the total body area (dotted line) and rankl-CFP*^+^* area in SKLT-treated (**A**) and control (**B**) fish. (**C**) Frequency distribution of the CFP mean intensity. (**D-E**) Quantifications of the mineralized area of the vertebral column in the different fish clusters. (**F**) Representative images depicting AR-S stained vertebral column of control (upper image) and SKLT-exposed (lower image) fish in High CFP intensity group. (**G**) Representative images depicting the count of mineralized neural arches in control (upper image) and SKLT-exposed (lower image) fish in High CFP intensity group. (**H**) Number of mineralized neural arches in SKLT-exposed and control fish in High CFP intensity group. Statistical differences were tested through an unpaired *t* test (*p*<0.05). Asterisks indicate values statistically different with *p*<0.01 (**). Scale bars are 360 µm in **A** and **B**, and 730 µm in **J** and **K**.

### SKLT inhibits osteoclastic differentiation in murine macrophages

The capacity of SKLT to inhibit osteoclastic differentiation was assessed in the murine macrophage cell line RAW 264.7, which differentiates into active osteoclasts upon treatment with RANKL. First, we exposed cells to different concentrations of SKLT to evaluate the effect of the extract on cell survival and proliferation. While no cytotoxicity was observed in RAW 264.7 cells exposed for 24 and 48 h to SKLT at the concentrations of 50, 100, and 200 µg/mL (**Figure 7A**), cell proliferation was significantly reduced at 1, 3, 5, and 7 days of exposure, even when cells were exposed to the lowest concentration of SKLT at 50 µg/mL (**Figure 7B**). Subsequently, cells were co-exposed for 6 days to RANKL at 50 ng/mL and SKLT at 50 µg/mL, and osteoclast differentiation was assessed through DAPI and TRAP stainings (**Figure 7C**, **D**). While no large multinucleated (> 2 nuclei) TRAP+ cells, i.e. osteoclasts, were observed in the negative control group (no RANKL), 14.3 ± 2.6 osteoclasts/5x field were counted in the RANKL-treated group.

**Figure 7.**
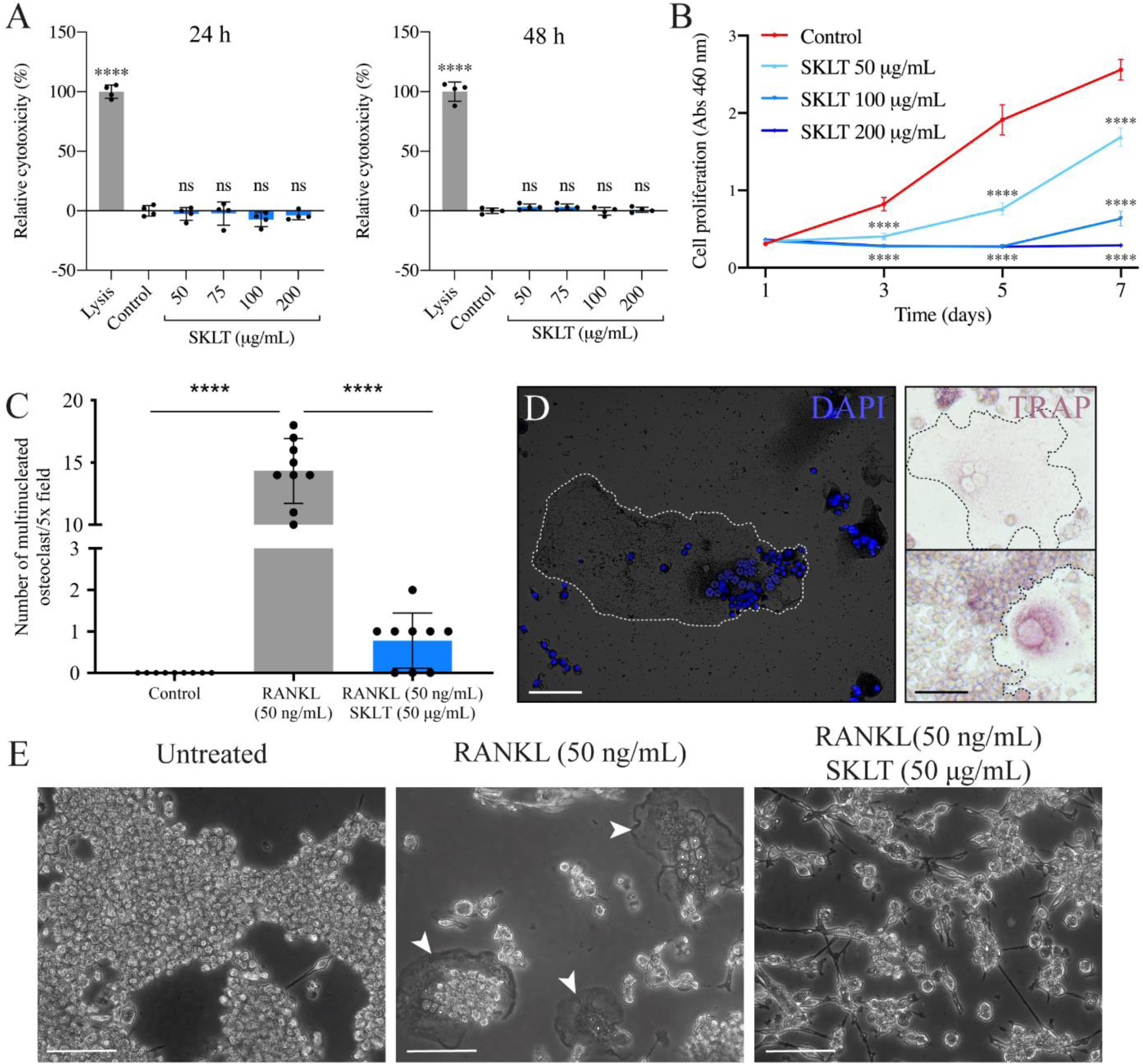
SKLT inhibits cell proliferation and osteoclastic differentiation in murine macrophages. Cytotoxicity, proliferation and osteoclastic differentiation of RAW 264.7 cells were assessed upon exposure to the ethanolic extract of *S. costatum* (SKLT) or 0.1% ethanol (Control). (**A**) Cytotoxicity (LDH assay) in cells exposed for 24 h (left) and 48 h (right) to different concentrations of SKLT. (**B**) Proliferation (XTT assay) of cells exposed to to different concentrations of SKLT. (**C**) Number of multinucleated osteoclasts per field (magnification 5.0X). (**D**) Representative fluorescence image of DAPI stained multinucleated osteoclasts (left) and brightfield images of TRAP^+^ osteoclasts (right). (**E**) Representative phase-contrast images of the 3 experimental groups. Arrowheads indicate multinucleated osteoclasts. Differences between each treatment and the control were tested though a one-way ANOVA followed by Dunnett’s multiple comparison test (*p*<0.05). Asterisks indicate values statistically different with *p*<0.0001 (****). Scale bars are 150 µm.

Co-treatment with SKLT significantly reduced the number of differentiated osteoclasts to 0.78 ± 0.67 osteoclast/5.0x field (**Figure 7C**, **E**). These data highlight the anti-proliferative and anti-osteoclastogenic effects of SKLT on mouse macrophages.

## Discussion

The immune system is increasingly recognized as a primary driver of several bone erosive pathologies^53^. For example, in primary osteoporosis, a complex dysregulation of both innate and adaptive immune cells contribute to establish chronically elevated bone resorption levels^30^.

Although no therapeutics currently available act through immunomodulatory mechanisms, recent findings supporting the roles of different immune cells in modulating bone homeostasis have provided the molecular basis for the exploration of immunomodulatory therapies to treat diseases like osteoporosis^54^.

In this work, we aimed at providing molecular insights on the potential of a novel immunomodulatory approach for treating bone loss, by implementing an emerging natural extract from the microalgae *Skeletonema costatum*.

In a zebrafish model of fin regeneration^44^, this extract shifted the mineral equilibrium of regenerating rays towards pro-mineralogenic outcomes. In detail, exposure to SKLT inhibited ray bifurcation, resulting in regenerated rays with little to no bifurcation. Previous studies have elucidated the complexity and tight regulation of the process of *de novo* ray bifurcation during fin regeneration. In detail, ray branching is controlled by a dynamic interaction of de-differentiated and subsequently re-differentiated osteoblastic cells^55–57^, and elongated osteoclastic cells named osteolytic tubules (OLTs), which exert an anti-stitching action over the advancing mineralization front, precisely determining the position of the bifurcation point along the ray axis^47^.

The same study showed that exposure to pro-osteogenic or anti-resorptive pharmaceuticals could modulate the proximo-distal positioning of the bifurcation point, making the evaluation of proximalization/distalization of ray bifurcation a suitable screening system for pro-osteogenic or anti-osteoclastogenic activities^47^.

Thus, the distalization of ray bifurcation in regenerating fins exposed to SKLT may result from compounds that enhance bone deposition or inhibit bone resorption.

In light of recent studies reporting an anti-inflammatory activity in *S. costatum* extracts *in vitro*^42,43^, and highlighting the importance of inflammation for osteoclastic differentiation^16^, we hypothesized that anti-inflammatory and immunomodulatory compounds in SKLT inhibited osteoclast function. This hypothesis was validated by a reduced presence of cells expressing cathepsin k – a marker of osteoclast precursors – and a weaker activity of the tartrate-resistant acid phosphatase (TRAP) – a marker of mature osteoclast function, consistent with data previously reported^47^.

We, therefore, hypothesized that SKLT contains compounds that can delay the recruitment and/or differentiation of osteoclast precursors during the early stages of caudal fin regeneration.

A reduction of fin regenerative performances, demonstrated by a decrease in the area of the regenerative outgrow at the late stages of fin regeneration, was also observed in fish exposed to SKLT. We propose that this effect may result from the immunomodulatory action of SKLT. Indeed, fin regeneration is characterized by a quick response in which immune cells invade the wound and release pro-inflammatory cytokines^49,58^. This results in an inflammatory flare, which is not only a consistent event in all regenerative processes across different organs, including the fin^49^, heart^59,60^, and spinal cord^61^, but also a necessary step enabling and somehow triggering the activation of pro-regenerative molecular programs in regenerative species like zebrafish^62^.

In this regard, it has been shown that zebrafish larvae with pharmacologically suppressed inflammation exhibit a reduced regenerative capacity, i.e. the regenerated area of the larval fin fold is reduced^63,64^. Thus, we believe that the reduced regenerative capacity of fish exposed to SKLT is related to its anti-inflammatory activity. Similar results were observed in adult zebrafish, where genetic ablation of macrophages not only reduced regenerative performances, but also inhibited the bifurcation of bony rays^65^.

Insights into these notable effects – i.e. impaired fin regeneration and altered fin ray mineralization and patterning – gained from the transcriptomic analysis of caudal fin blastemas exposed to SKLT revealed a downregulation of genes typically associated with a wound-induced inflammation. Among those, interleukin 20 receptor alpha (*il20ra*) was of particular interest, as IL20 is a pro-inflammatory and pro-osteoclastogenic cytokine involved in various osteoerosive mechanisms, including estrogen deficiency-induced bone loss^66^, and orthodontic alveolar bone remodeling^67^.

Adaptive immunity was also affected by SKLT as genes related to T-cell activation, interferon γ modulation, and antigen presentation were downregulated in animals exposed to *S. costatum* extract. The notion that estrogen deficiency leads to increased T-cell activation, comes from murine studies in the early 2000s^68^ when the term osteoimmunology was first coined by Arron & Choi^69^. More than 20 years of studies have helped clarify the role of specific subfamilies of T-lymphocytes as major contributors to bone loss in different erosive pathologies^30^. Here, several genes involved in T-cell differentiation and activation, such as *cd4*, *cd74a*, *cd74b*, *ccl19a.1*, and *jak1* were downregulated in SKLT-exposed fish, showing reduced osteoclastic differentiation.

Various components of the interferon (*ifng1r*, *irf4b*, *irf10*), which regulate the differentiation and activity of different T-cell subtypes^70–72^, were here downregulated by SKLT exposure. In addition, *itk*, a gene required for the differentiation of natural killer T cells^73^, and the IL-7 receptor (*il7ra*), associated with the production of pro-osteoclastogenic cytokines^74^, and uncoupling of bone resorption and bone formation^75^, were downregulated. Interleukin 13 (IL-13), typically produced by anti-inflammatory Th2 lymphocytes^76^, is considered an osteoprotective cytokine, inhibiting osteoclastogenesis^77^. Here, fish exposed to SKLT displayed an increased expression of the IL13 receptor (*il13ra2*).

Interestingly, SKLT also modulated genes related to cell-cell communication, some of which were recently associated with the paracrine communication between osteoblasts and macrophages during their recruitment and differentiation into osteoclasts. In this context, the chemokine Cxcl9l (orthologous to human CXCL9) is produced in medaka under osteoporotic conditions by pre-osteoblasts and guides the fate of a subset of macrophages expressing the chemokine receptor Cxcr3.2 (orthologous to human receptor CXCR378) toward osteoclastic differentiation^78^.

In zebrafish, the chemokine receptor Cxcr3 is coded by 3 genes – *cxcr3.1*, *cxcr3.2* and *cxcr3.3*. Here, *cxcr3.1* was downregulated in fish exposed to SKLT, possibly indicating a reduction of a specific population of macrophages, highly responsive to pro-osteoclastogenic signaling. Another osteoblast-secreted paracrine factor that recently emerged as important for osteoclastic function is the matrix-metalloproteinase 13^79^. The analysis of medaka transcriptome in osteoporotic conditions revealed that osteoblast progenitors strongly express *mmp13b*, and that *mmp13b* loss-of-function resulted in osteoclasts being immature, inactivated, and unable to resorb the extracellular matrix^79^. In the present work, fish exposed to SKLT had a reduced expression of *mmp13b*.

Altogether, these data demonstrate that SKLT induced a complex modulation of the immune response in 24 hours post-amputation blastemas, with a marked suppression of molecular programs associated with antigen presentation, T-cell differentiation and activation, and altered paracrine communication responsible for the recruitment of macrophages to the bone niche, and their differentiation into osteoclastic cells.

A medaka model of induced osteoporosis, where the systemic overexpression of RANKL leads to extensive bone loss^19^, was used to confirm that SKLT immunomodulatory effect can protect against bone erosion when it is caused by a pathologically elevated osteoclast formation and activation. In combination with our molecular data on zebrafish, these findings suggest that SKLT, by modulating inflammation and T cell differentiation, efficiently reduced bone loss in a disease setting. Such inhibition could be the result of a reduction in the differentiation of macrophages into the osteoclastic lineage.

The translational applicability of these results to a mammalian system was confirmed *in vitro* using a murine macrophages cell line. A clear anti-proliferative effect, coherent with the results obtained in zebrafish, and a reduced formation of multinucleated osteoclasts was observed in macrophages exposed to RANKL.

Altogether, our data evidences the therapeutic potential of molecules that modulate both the innate and adaptive immune responses for bone erosive pathologies, particularly T-cell activation, antigen presentation, and macrophage fate determination to osteoclastogenesis. We revealed the presence of compounds with immunomodulatory and anti-osteoclastogenic bioactivities in *Skeletonema costatum*, opening the possibility for a possible future implementation of a treatment of bone erosive disorders. Whether such biological activities result from the action of a single compound, or the synergistic effect of different molecules was not addressed here; but it is a question that needs further examination to validate the pharmacological applicability of this natural extract. In this regard, it is worth mentioning that various authors have argued in favor of a more holistic approach for the treatment of chronic diseases displaying bone loss, with the rationale that natural nutraceuticals with broad-spectrum immunomodulatory properties could offer an alternative path to address the therapeutic needs of a diverse set of bone erosive disorders^80–82^. Yet, in the present scenario, the identification of such compounds must be a priority for further exploitation of this microalga for pharmacological applications.

Notwithstanding, our study provide a proof of concept for the osteoprotective potential of compounds that modulate immune cells and inflammation.

## Methods

### Preparation of microalgae extracts

Freeze-dried biomass of *Skeletonema costatum* (Necton S.A., Olhão, Portugal) was macerated with 96% ethanol (Laborspirit Lda, Lisbon, Portugal) using a biomass-solvent ratio of 1 g:40 mL (M/V), by gently stirring at 24 °C for 18 h. The macerate was centrifuged for 5 min at 1,000 ˟ g using an Allegra 6R centrifuge (Beckman Coulter Inc, Brea, USA) and the supernatant was collected. The pellet was washed twice with 96% ethanol and all supernatants were pooled and then vacuum filtered sequentially through 0.45 µm and 0.22 µm nylon membranes (Labbox Labware S.L., Barcelona, Spain). Filtrate was concentrated with a rotatory evaporator RV 10 digital (IKA-Werke GmbH & Co. KG, Staufen im Breisgau, Germany), with the temperature set at 40 °C and pressure at 178 mbar, until obtaining a dense, paste-like extract. Extraction yield – 37.9 ± 2.4% – was calculated from 2 mL aliquots (n = 3) placed under a gentle flow of 99.8% nitrogen until complete evaporation of the solvent.

### Fish maintenance

Zebrafish wild type line AB and transgenic line Tg(Ola.*ctsk*:FRT-DsRed-FRT-Cre,*myl7*:EGFP)^mh201^, 48, referred to as Tg(*ctsk*:DsRed) throughout the manuscript, were maintained in a water recirculating system ZebTEC (Tecniplast, Buguggiate, Italy) with the following conditions: temperature 28 ± 0.1 °C, pH 7.5 ± 0.1, conductivity 700 ± 50 μS, ammonia and nitrites at levels below 0.1 mg/L, nitrates lower than 50 mg/L, and a photoperiod of 14:10 h light-dark. Medaka transgenic line Tg(*rankl*:HSE:CFP)^TG1135^, hereafter referred to as Tg(*rankl*:HSE:CFP), were purchased from the National BioResource Project Medaka (NBRP Medaka)^83^ and maintained in a water recirculating system with the following conditions: temperature 27 ± 0.1 °C, pH 7.0 ± 0.1, conductivity 300 ± 100 μS, ammonia and nitrites at levels below 0.1 mg/L, nitrates lower than 50 mg/L and a photoperiod of 14:10 h light-dark. For zebrafish and medaka, system water was prepared by supplementing reverse osmosis water with a salt mixture (Instant Ocean, City, USA) and sodium bicarbonate (Sigma-Aldrich, St. Louis, USA). All fish were fed daily with the commercial dry food Zebrafeed (Sparos Lda, Olhão, Portugal).

### Zebrafish caudal fin regeneration assay

The caudal fin of wild-type or transgenic adult zebrafish aged 3-4 months was amputated 1-2 segments anterior to the bifurcation of the most peripheral branching lepidotrichia, as described by Cardeira et al. (2016)^44^. After finectomy, fish (n > 14) were placed at 33 ± 1 °C in 3 L-plastic containers at a density of 5 fish/L. Fish were exposed for 5 days (AB wild-type) for the first experiment evaluating the mineralogenic performances, and for 10 days (Tg(*ctsk*:DsRed)) for the experiments tracking osteoclastic cells, to the *S. costatum* ethanolic extract (SKLT) at 56 µg/mL or to ethanol (vehicle) at 0.1%, supplemented in system water. Treatment was renewed daily. Moderate water dynamics and air-water exchanges were facilitated by bubbling fish tanks with an air pump. Water quality parameters were monitored daily and maintained stable for the duration of the experiment as follows: dissolved oxygen 7.0 ± 0.5 mg/L, pH 7.4 ± 0.2 and conductivity 680 ± 20 μS. Wild-type fish were sacrificed at 120 hours post-amputation (hpa) using lethal anesthesia of 0.6 mM tricaine methanesulfonate (MS-222, pH 7.0; Sigma-Aldrich), then immersed for 30 min in 0.03% alizarin red S (AR-S, pH 7.4; Sigma-Aldrich) and washed two times for 5 min with system water. Stained fish were imaged for fin morphometric analysis. Transgenic fish Tg(*ctsk*:DsRed) were sampled (n ≥ 9) at different time points (24, 48, 72, 96, 120 and 240 hpa), then immersed for 30 min in 0.2% calcein (pH 7.4; fluorexon, Sigma-Aldrich), washed twice in system water for 10 min. After the staining, fish were anesthetized for 5 min in tricaine and imaged. For Bulk RNAseq analysis, fin blastemas were collected at 24 hpa, pooled (n = 5, 10 blastemas per pool), and conserved at -80 °C until further processing.

### TRAP activity in caudal fins

Tg(*ctsk*:DsRed) zebrafish (n ≥ 8) were sacrificed at 24 and 240 hpa with a lethal dose of tricaine (for detail, see previous section). Caudal fins were amputated at the level of the caudal peduncle, washed once with 1X phosphate buffer saline (PBS, pH 7.4) and fixed for 4 h in 4% paraformaldehyde solution (PFA, solubilized in PBS, pH 7.4) at 24 °C. Tartrate resistant acid-phosphatase (TRAP) staining was performed as previously described by Blum & Begemann (2015)^52^ and fins were imaged as described below.

### Morphometric analysis of caudal fins

Fins were imaged under an MZ10F fluorescence stereomicroscope (Leica, Wetzlar, Germany) coupled to a DFC7000T color camera (Leica). Bright-field images were collected to assess the progression of fin regeneration. Fluorescence images were collected to assess (i) *de novo* bone formation in wild type fish stained with AR-S or calcein, and (ii) the involvement of *ctsk*-expressing cells in transgenic fish labelled with DsRed. Bright-field images were acquired with an exposure time of 1 ms. Fluorescence images were acquired with the filter set ET560/40x - ET630/75m and an exposure time of 600 ms for mCherry, and the filter set ET470/40x - ET525/50m and an exposure time of 80 ms for GFP. Other image parameters were: gamma 1.00; image format 1920×1440 pixels; and binning 1×1. Fluorescence images were analyzed using ImageJ software version 2.0.0-rc-69/1.52p and processed using the ZFBONE toolset for caudal fin morphometrics^84^. Fin regeneration and mineralization were assessed following the method described by Cardeira et al. (2016)^2^, by calculating the regenerated area (REG), the stump width (STU), the mineralized area (MIN), and the average width of the rays before the amputation (RAYs). Fin ray patterning was also assessed by calculating the average ray width ratio 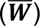 and the average bifurcation ratio 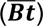 as shown in **Supplementary equations a** and **b**. For the quantification of *ctsk* signal in transgenic fish, DsRed-positive area was measured using a color threshold on fluorescent images and normalized with REG/STU. To quantify the TRAP signal, TRAP-positive areas were measured using a color threshold on bright-field images, and subsequently normalized with REG/STU.

### RNA preparation

Total RNA was extracted from pools of blastemas at 24 hpa (n = 5) using NZYol (NZYTech, Lisbon, Portugal) and quantified using a NanoDrop OneC spectrophotometer (Thermo Fisher Scientific, Waltham, USA). RNA integrity was confirmed using an Experion Automated Electrophoresis system (Bio-Rad, Hercules, USA). Only RNA with an RNA integrity number (RIN) higher than 7 was used.

### Bulk RNA sequencing and analysis of differentially expressed genes

Bulk RNA sequencing was outsourced to STABVIDA Lda (Caparica, Portugal). DNA libraries were constructed using a Stranded mRNA Library Preparation kit (STABVIDA) and sequenced on a Novaseq platform (Illumina, San Diego, USA) to generate 150 bp paired-end sequencing reads. Raw sequence data was processed using CLC Genomics Workbench 12.0.3^85^. Trimming was done in 3 steps: Quality trimming based on quality scores (error probability 0.01); Ambiguity trimming (ambiguous limit of 2 nucleotides); and Length trimming to discard reads shorter than 30 nucleotides. The quality-checked sequencing reads were mapped against zebrafish reference genome GRCz11 (GCF_000002035.6) using length fraction and similarity fraction equal to 0.8. TPM (transcripts per million) and RPKM (reads per kilobase of transcript per million mapped reads) were then determined from mapped data^86^. Differential expression analysis was performed with the multi-factorial EdgeR method in R^87^. Overall gene expression results were represented with a clustering heatmap of the gene expressions, with a principal component analysis (PCA) plotting the amount of variance explained by the two principal components and a volcano plot representing overall gene expression with fold change in the x-axis and significance of expression in the y-axis (**Supplementary Figure S2**). These overall representations of gene expression were performed to assess variation patterns in the gene expression dataset and identify outlier samples for quality control. For functional annotation enrichment analysis, the lists of upregulated and downregulated genes were analyzed with the online resource Database for Annotation, Visualization and Integrated Discovery (DAVID)^88,89^ for Gene Ontology – Biological Processes (GO:BP) with FDR<0.01 and enriched biological processes.

### Medaka model of RANKL overexpression-induced osteoporosis

Eggs of the medaka line Tg(*rankl*:HSE:CFP) were produced following an in-house breeding program and maintained in Petri dishes with 40 mL of system water supplemented with 0.0002% (w/v) methylene blue until 8 days post-fertilization (dpf). At 8 dpf, hatched larvae were placed at 39 °C for 2 h (heat-shock) and screened for CFP (cyan fluorescent protein) signal using a fluorescence microscope (see parameters below). CFP-positive fish were distributed into a 6-well plate at the density of 5 larvae/well. Each well was filled with 10 mL of system water supplemented with SKLT at 56 µg/mL or 0.1 % ethanol (vehicle). Treatment was renewed 100% daily. Six days after heat-shock (dahs), larvae were sacrificed with a lethal dose of tricaine (see above), stained with AR-S and imaged as described above. Bright-field and fluorescence images were collected for each fish. Bright-field images were acquired using an exposure time of 2 ms. AR-S fluorescence images were acquired using the filter set ET560/40x - ET630/75m and an exposure time of 700 ms. CFP fluorescence images were acquired using the filter set ET436/20x - ET480/40m and an exposure time of 200 ms. Other image parameters were: gamma 1.00, image format 1920×1440 pixels, and binning 1×1. Images were analyzed using ImageJ using two ad hoc macros (**Supplementary macro 1** and **2**). CFP fluorescence images were transformed to 8-bit images, and the CFP-positive area was measured using a color threshold (min intensity 7 and max intensity 255). Pixel mean intensity inside the CFP-positive area was calculated and used as a proxy for CPF fluorescence intensity. AR-S fluorescence images were transformed to 8-bit images, and AR-S positive areas were measured using a color threshold (min intensity 5 and max intensity 255). AR-S positive area was normalized using the total body area (determined manually from bright-field images) to correct for inter-specimen size variation. AR-S positive area in abdominal and caudal vertebrae was used as a proxy of the mineralization of the vertebral column. The nomenclature proposed by Di Biagio et al. (2022)^90^ was used to identify vertebrae from 4 to 29. The number of mineralized neural arches was manually counted in each fish from AR-S images. Following the collection of the data, we determined the frequency distribution of CFP intensity and tested data normality through an Anderson-Darling test (p < 0.05). Normality was not met in the control group (p = 0.0035), but CFP intensity was normally distributed in SKLT-exposed fish (p = 0.220). Data distribution for CFP intensity was right-skewed (median < mean) for the control group and bimodal for the SKLT group. These findings implied that clustering was necessary to proceed with the statistical analysis. Fish were therefore grouped according to CFP intensity into “High CFP” (mean pixel intensity > 50) and “Low CFP” (mean pixel intensity < 50) for further analysis.

### Culture of mouse RAW 264.7 macrophages

RAW 264.7 cells were cultured in 10 mL cell culture dishes with Dulbecco’s Modified Eagle Medium (DMEM) supplemented with 10% fetal bovine serum (FBS, Sigma-Aldrich), 1% penicillin– streptomycin, 1% L-glutamine, and 0.2% fungizone at 37 °C in a 5% CO2-humidified atmosphere. Pre-confluent cell cultures were sub-cultured 1:4 every other day using trypsin-EDTA solution (0.2% trypsin, 1.1 mM EDTA, pH 7.4). All cell culture reagents were from GIBCO-ThermoFisher Scientific, unless otherwise stated.

### Cytotoxicity and cell proliferation

LDH Cytotoxicity Assay kit and XTT Cell Proliferation Assay kit (Canvax Biotech, Córdoba, Spain) were used, respectively, to evaluate the effects of SKLT extracts on cellular toxicity and proliferation. For cytotoxicity, RAW 264.7 cells were seeded in a 96-well plate at 1.0 × 104 cells/well (n = 4) in 100 µL of culture medium supplemented with either the SKLT extract at 50, 75, 100, 200 µg/mL or 0.1% ethanol (vehicle) and cultured for 24 and 48 h. Relative cytotoxicity was calculated as a percentage of LDH lysis control. For cell proliferation, RAW 264.7 cells were seeded in a 96-well plate at 500 cells/well (n = 6) in 100 µL of culture medium supplemented with either the SKLT extract at 50, 75, 100, 200 µg/mL or 0.1% ethanol (vehicle) and cultured for 1, 3, 5 or 7 days. Culture medium was renewed every other day. Relative cell proliferation was calculated as a percentage of the negative control.

### Osteoclast differentiation

RAW 264.7 cells were seeded in 12-well plates at 1.5 × 104 cells/well in 1 mL of culture medium (as described above) supplemented with either SKLT extract at 50 µg/ml or 0.1% ethanol (vehicle) and treated for 6 days with 50 ng/mL of RANKL (Preprotech, London, UK), prepared in 0.1% BSA, to induce their differentiation into multinucleated osteoclasts. Experimental conditions were: undifferentiated control (0.1% ethanol); differentiated control (50 ng/mL RANKL+ 0.1% ethanol); differentiated cells exposed to SKLT extract (50 ng/mL RANKL+ 50 µg/mL SKLT in 0.1% ethanol). The culture medium was freshly prepared and replaced daily. Osteoclast differentiation was assessed by counting the number of tartrate-resistant acid phosphatase (TRAP) positive cells and the number of nuclei following 4′,6-diamidino-2-phenylindole (DAPI) staining. For this, cells were fixed in 4% PFA at 24 °C (RT) for 10 min and stained with a protocol from Blum & Begemann (2015)^52^. Cells were subsequently stained with DAPI for nuclei detection and count. Cells were imaged using an Axio Vert.A1 inverted microscope (ZEISS, Jena, Germany) coupled with (i) an Axiocam 202 monocolor camera (ZEISS) for DAPI images, or (ii) a VWR VisiCam 5 Plus (VWR, Radnor, USA) for bright-field TRAP images. Number of multinucleated osteoclasts per field (5.0X magnification; n = 9) was calculated considering only TRAP+ cells with at least two nuclei.

### Statistical analysis

For all the experiments, normality was tested with a D’Agostino-Pearson omnibus normality test or with an Anderson-Darling test (p < 0.05). Homoscedasticity was tested through the Brown-Forsythe test (p < 0.05). When the distribution of the data of all the experimental groups resulted in normal and homogeneous, statistical differences between the control and the extract were tested with either an Unpaired t test or a one-way ANOVA followed by Dunnett’s multiple comparison test (p < 0.05). If the distribution of the data of any of the experimental conditions resulted in non-normal or nonhomogeneous, statistical differences between the control and the extract were tested with a Mann-Whitney test or a non-parametric test followed by Dunn’s multiple comparison test (p < 0.05). Statistical analyses were performed using Prism version 9.00 (GraphPad Software Inc., La Jolla, United States).

### Ethical statement

Procedures involving animals were performed following the EU and Portuguese legislation for animal experimentation and welfare (Directives 86/609/CEE and 2010/63/EU; Portaria 1005/92, 466/95 and 1131/97; Decreto-Lei 113/2013). Animal handling and experimentation were performed by qualified operators accredited by the Portuguese Direção-Geral de Alimentação e Veterinária under the authorization no. 012769/2021. All efforts were made to minimize pain, distress, and discomfort. Experiments were terminated (fish were returned to normal conditions or euthanized) whenever adverse effects were observed.

## Supporting information

Supplementary macros

## Acknowledgments

This work was financed by the European Maritime and Fisheries Fund (EMFF/FEAMP) through the National Operational Programme MAR2020 (grant 16-02-01-FMP-0057/OSTEOMAR), by the European Regional Development Fund (ERDF/FEDER) through the Transnational Cooperation Programme Atlantic Area (grant EAPA/151/2016/BLUEHUMAN), by the Marie Skłodowska-Curie innovative training network BIOMEDAQU (grant H2020-MSCA-ITN/766347), by National funds from the Portuguese Foundation for Science and Technology through projects UIDB/04326/2020 (DOI:10.54499/UIDB/04326/2020), UIDP/04326/2020 (DOI:10.54499/UIDP/04326/2020) and LA/P/0101/2020 (DOI:10.54499/LA/P/0101/2020), and through the doctoral fellowship 2021.05406.BD, and by the operational programmes CRESC Algarve 2020 and COMPETE 2020 through project EMBRC.PT ALG-01-0145-FEDER-02212. The authors would also like to thank João Navalho and Necton S.A. (Olhão, Portugal) for kindly providing us with the microalgal biomass used to prepare the ethanolic extracts evaluated in this work.

## Author Contributions

AC, MLC, VL, and PJG designed research. AC, KP, and JTR performed research. MT, HP contributed new reagents/analytic tools. AC, KP, SP, BL, analyzed data. AC, VL, PG wrote the paper. JTR, MLC, VL, PJG other (Provided supervision/founding acquisition/ revised the paper).

## Data availability statement

RNA sequencing data have been uploaded onto the free repository BioStudies EMBL-EBI with accession number E-MTAB-14677, and will be made publicly available upon article acceptance. All other raw data generated in this study will be made available upon request.

## Supplementary Material

**Figure S1.**
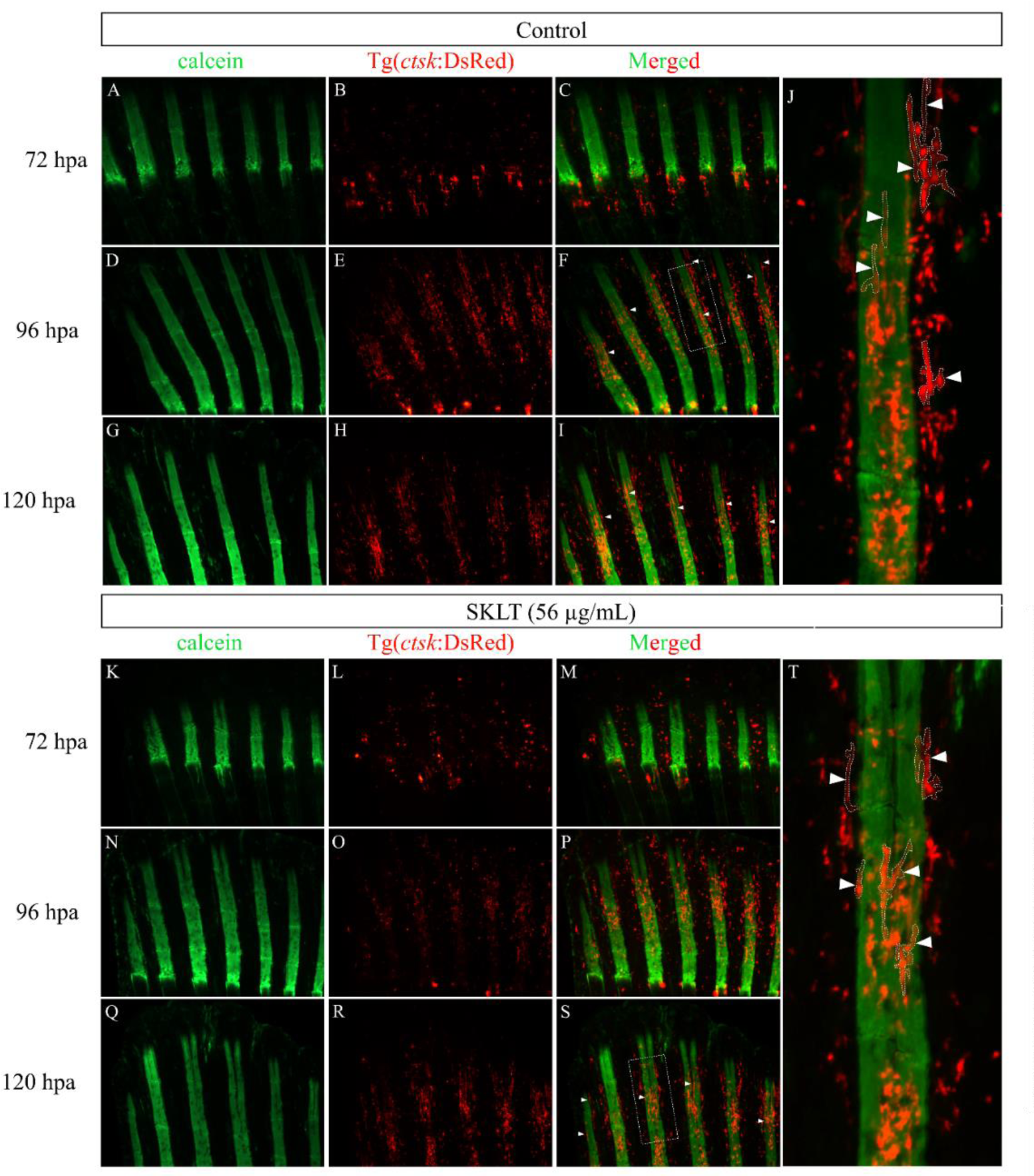
The ethanolic extract of *Skeletonema costatum* does not prevent the appearance of ctsk+ OLTs at later stages of fin regeneration. Presence of tubular and elongated *ctsk^+^* osteolytic tubules in the regenerating caudal fin of adult zebrafish exposed to 0.1% ethanol (Control; **A-J**) or to the ethanolic extract of *Skeletonema costatum* at the concentration of 56 µg/mL (SKLT; **K-T**). Osteolytic tubules (OLTs) are observed in most fins at 96 hpa in control fish (**J**), and at 120 hpa (**T**) in SKLT-treated fish. hpa, hours post amputation; white arrowheads, ctsk+ osteolytic tubules associated with the regenerated bony rays.

**Figure S2.**
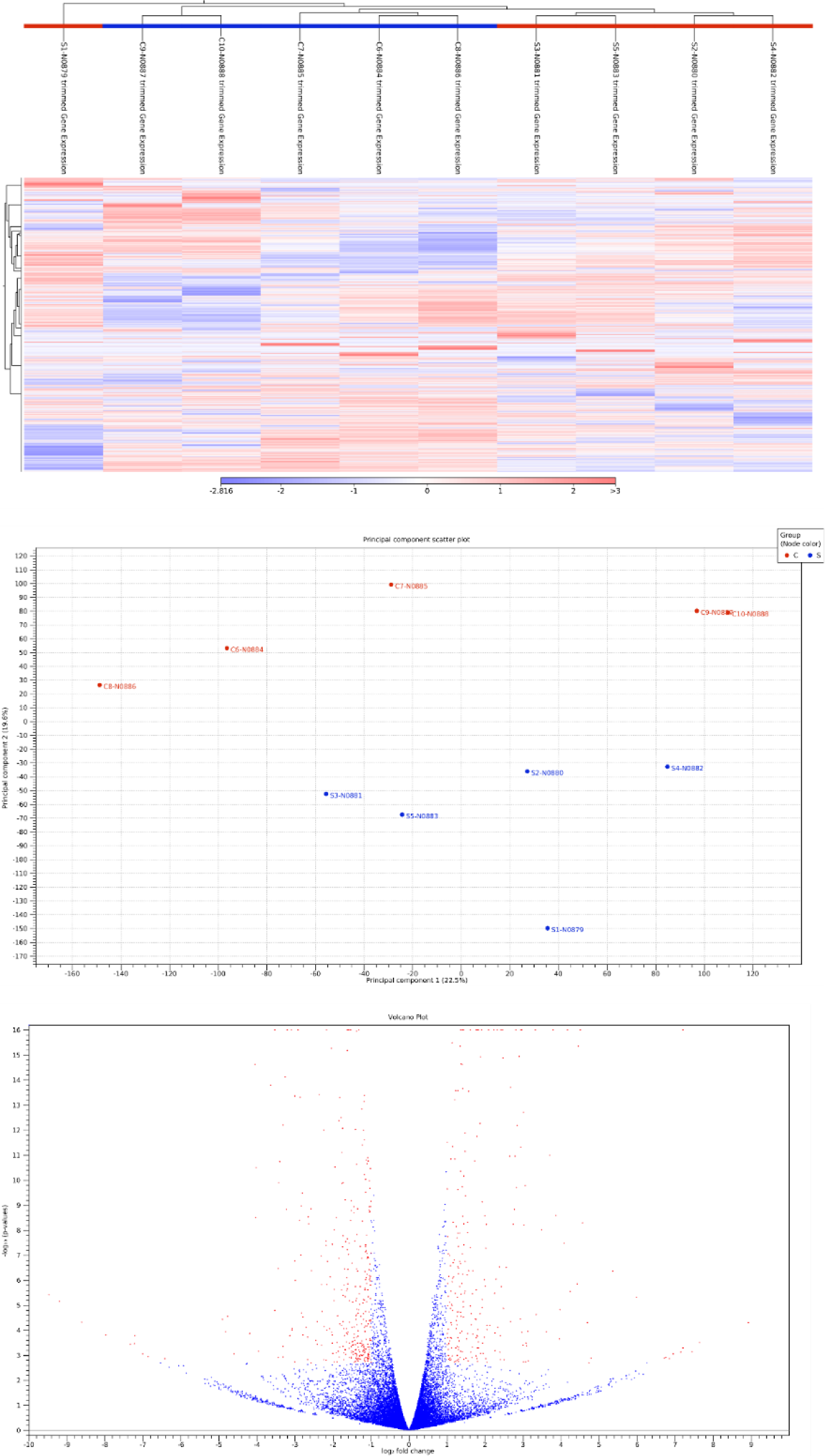
Overall gene expression results from RNAseq analysis of SKLT-exposed vs control specimens. (**A**) Heatmap of the expression analysis created with the results of the gene expression. The red colour represents over-expression, while the blue colour is under-expression. (**B**) PCA corresponds to the gene expression of the groups of samples. The first principal component is shown on the X-axis, and the second principal component is shown on the Y-axis. The value after the principal component identifier displays the amount of variance explained by this principal component. (**C**) Volcano plot representing overall gene expression with fold change in the x-axis and significance of expression in the y-axis, with significantly differentially expressed genes represented in red dots and not significantly differentially expressed genes in blue dots.

## Supplementary Equations

***(a)*** Ray width ratio 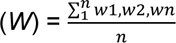, Average width ratio 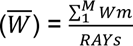 where,

*w* = width of the segments after the amputation

*n* = number of segments after the amputation

*RAYs* = average width of the first segment before the amputation

*m* = number of rays

***(b)*** Ray bifurcation ratio (*Bt*) = *BIF/TOT*, Average width ratio 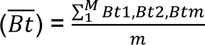 where,

*BIF* = bifurcation length

*TOT* = regenerated ray length

*m* = numbers of rays

