## Supplementary macros for "Harnessing the immune system to treat bone loss: The immunomodulatory and osteoprotective effects of the microalga *Skeletonema costatum*": Supplementary macro 1.ijm

```
/*             
DEVELOPER: Marco Tarasco                                     
EMAIL:                            
*/
//Macro developed for the analysis of body area in transgenic medaka.

macro "Measure area Action Tool - C000D06D09D15D1aD25D2aD34D37D38D3bD44D4bD53D54D5bD5cD63D6cD73D76D79D7cD83D87D88D8cD93D9cDa2DadDb2DbdDc1DceDd1Dd2Dd3DdcDddDdeDe0De1De4DebDeeDefDf4Dfb"{

//Menu options
	
	Dialog.create("Measure area");
	Dialog.addRadioButtonGroup("Mode", newArray("Single","Batch"),1, 2, "Single");
	Dialog.show();
	mode=Dialog.getRadioButton();
	
// analyse single image or batch?
				if (mode=="Single") {
					doAnalysis();
				}
				else {
					dir=getDirectory("Choose a Directory");
					list=getFileList(dir);
				 
							for (i = 0; i < list.length; i++) {
								if (endsWith(toLowerCase(list[i]), ".bmp")||endsWith(toLowerCase(list[i]), ".tif")||endsWith(toLowerCase(list[i]), ".jpg")){
									if(!File.exists(dir+"analyzed/"+list[i]+"_analyzed.jpg")){
						           		open(dir+list[i]);
						           		doAnalysis();
						           		run("Close All");
									}
								}   
							}
				}
		
function doAnalysis(){
	
	if (nImages==0) waitForUser("Please open an image");
	if (nImages==0) exit("No image was opened!!");
	run("Set Scale...", "distance=0 known=0 pixel=1 unit=pixel");
	roiManager("reset");
	roiManager("Show None");
	dir=getDirectory("image");
	img=getTitle();
	
//measure fish area
	setTool("polygon");
		while(selectionType==-1){
			waitForUser("Select "+"Area");
		} 
	roiManager("Add"); 
	List.setMeasurements;
	areaFish=List.getValue("Area");
	setResult("Label", nResults, img);
	setResult("Fish area", nResults-1, areaFish);
	roiManager("Select", 0);
	roiManager("Set Color", "magenta");
	roiManager("Rename", "Fish area");
	roiManager("Deselect");
	roiManager("show all without labels");
	run("Flatten");
	getDateAndTime(year, month, dayOfWeek, dayOfMonth, hour, minute, second, msec);
	dateTag=""+year+"-"+IJ.pad((month+1), 2)+"-"+IJ.pad((dayOfMonth), 2)+"_";
	direct=dir+"analyzed/";
	File.makeDirectory(direct);
	roiManager("Save",direct+img+"_RoiSet.zip");
	saveAs("jpeg",direct+img+"_analyzed");
	Table.save(direct+dateTag+"_Results.xls");
	run("Close All");
}
}
```
